# A neural model for insect steering applied to olfaction and path integration

**DOI:** 10.1101/2020.08.25.266247

**Authors:** Andrea Adden, Terrence C. Stewart, Barbara Webb, Stanley Heinze

## Abstract

Many animal behaviours require orientation and steering with respect to the environment. For insects, a key brain area involved in spatial orientation and navigation is the central complex. Activity in this neural circuit has been shown to track the insect’s current heading relative to its environment, and has also been proposed to be the substrate of path integration. However, it remains unclear how the output of the central complex is integrated into motor commands. Central complex output neurons project to the lateral accessory lobes (LAL), from which descending neurons project to thoracic motor centres. Here, we present a computational model of a simple neural network that has been described anatomically and physiologically in the LALs of male silkworm moths, in the context of odour-mediated steering. We present and analyze two versions of this network, both implemented in the Nengo framework, one rate-based and one based on spiking neurons. The modelled network consists of an inhibitory local interneuron and a bistable descending neuron (‘flip-flop’), which both receive input in the LAL. The flip-flop neuron projects onto neck motor neurons to induce steering. We show that this simple computational model not only replicates the basic parameters of male silkworm moth behaviour in a simulated odour plume, but can also take input from a computational model of path integration in the central complex and use it to steer back to a point of origin. Furthermore, we find that increasing the level of detail within the model improves the realism of the model’s behaviour. Our results suggest that descending neurons originating in the lateral accessory lobes, such as flip-flop neurons, are sufficient to mediate multiple steering behaviours. This study is therefore a first step to close the gap between orientation circuits in the central complex and downstream motor centres.

**Author summary:** Targeted movements and steering within an environment are essential for many behaviours. In insects, the brain’s center for spatial orientation and navigation is the central complex, which processes information about the configuration of the local environment as well as global orientation cues such as the Sun position. Neural networks in the central complex also compute the insect’s heading direction, and are thought to be involved in generating steering commands. However, it is unclear how these steering commands are transmitted to downstream motor centers. Output neurons from the central complex project to the lateral accessory lobes, a neuropil which also gives rise to descending pre-motor neurons that are involved in steering in the silkworm moth Bombyx mori. In this study, we provide a computational model of a pre-motor neural network in the lateral accessory lobes. We show that this network can steer an agent towards the source of a simulated odor plume, but that it can also steer efficiently when getting input from an anatomically constrained network model of the central complex. This model is therefore a first step to close the gap between the central complex and thoracic motor circuits.

## Introduction

Insects display an astonishing range of behaviours that include highly directed movements. For example, male moths navigate towards females emitting pheromones [1] [2] and female crickets move towards singing males [3] [4]. Other insects use visual cues to maintain a straight heading over short or long distances (dung beetle: [5] [6]; Monarch butterfly: [7]; Bogong moth: [8]), and can even rely purely on memory to navigate home [9] [10]. While the cues used for navigation are different in these examples, they elicit very similar behaviours: upon encountering an appropriate stimulus, the animal chooses a direction with respect to that stimulus and begins moving in that direction. If the stimulus is temporarily lost, searching behaviour is initiated. Thus, the motor patterns elicited by different kinds of stimuli can be remarkably similar.

The neural mechanisms underlying some of these behaviours are well-known, while they are less well described for others. However, it is clear that the spatial context for orientation and navigation is computed in the central complex (CX), which is the only unpaired and midline-spanning neuropil in the insect brain (Fig. 1; [11]). In recent years, progress has been made in modelling this ‘compass system’ of insects, showing that part of the CX network can be modelled as a ring attractor. A ring attractor can be thought of as a ring-shaped neural network, in which an activity ‘bump’ exists in one location, while all other locations on the ring are inhibited. The activity bump can then be moved around the ring by external inputs as well as self-generated angular velocity cues. The position of the bump in the CX ring attractor corresponds to the heading direction of the animal in its environment, thus providing a reliable internal representation of the animal’s heading [12] [13]. An extended model of the CX network furthermore showed that the CX is a possible substrate for the path integrator [14], i.e., continuously integrating velocity to maintain an estimate of the direction and distance to a reference location. This model also demonstrates how CX output can serve directly as a steering command: the summed activity of columnar output neurons in each hemisphere is compared and any imbalance between the two hemispheres should produce a turn towards the relevant side, while a balanced output results in straight movement.

**Fig 1.**
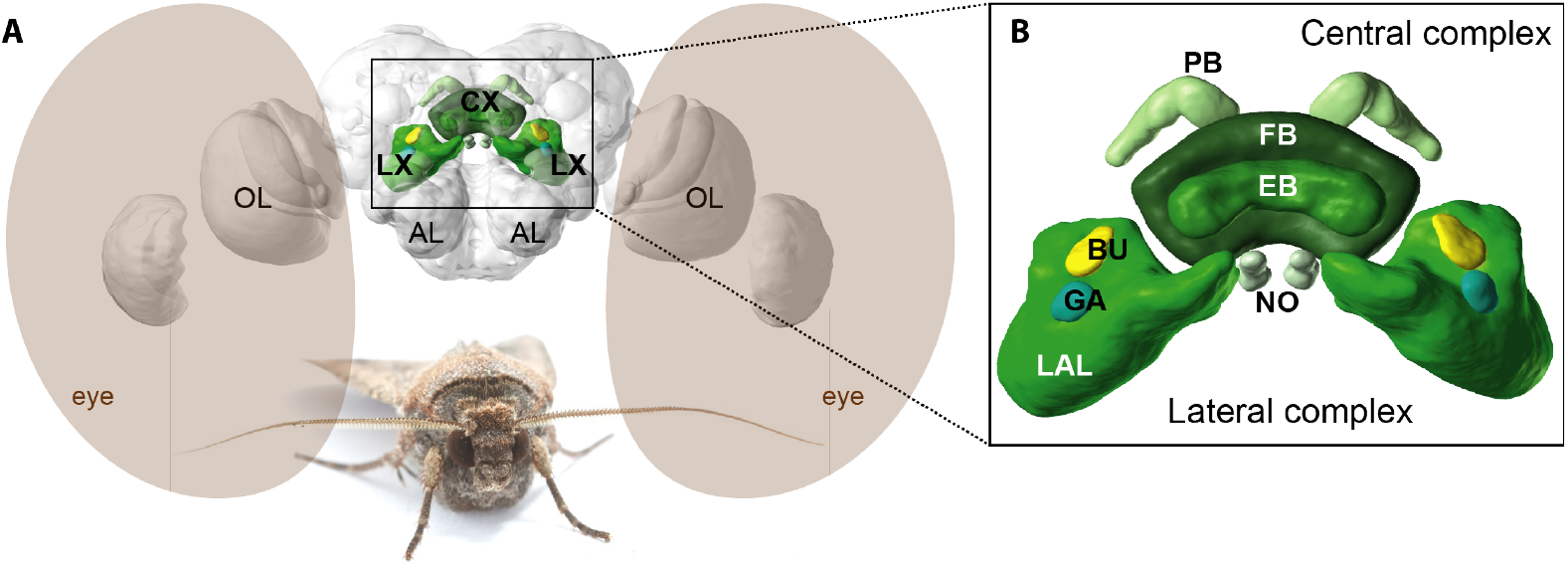
Insect brain and central complex / lateral complex anatomy. A: The central complex (CX) is located in the center of the insect brain and is the only midline-spanning neuropil. It is flanked by the lateral complexes (LX). Also marked are the optic lobes (OL) and the antennal lobes (AL). B: The CX consists of the protocerebral bridge (PB), the fan-shaped body (FB), the ellipsoid body (EB) and the paired noduli (NO). The LX consists of the lateral accessory lobes (LAL), the bulb (BU) and the gall (GA). CX output neurons project to the LAL. The brain shown is a Bogong moth brain (Agrotis infusa [31] [32]) retrieved from insectbraindb.org, species handle https://hdl.handle.net/20.500.12158/SIN-0000002.1. Photo courtesy of Ajay Narendra.

Despite our increasingly complete understanding of these networks, a question that remains unanswered is how the CX output is actually translated into motor control; i.e. how it might be integrated in thoracic motor centres to influence behavioural decisions. CX output neurons project to the lateral accessory lobes (LAL; figure 1), a brain region that has been described as a pre-motor centre, because several types of descending neurons that project to thoracic motor centres have post-synaptic endings in the LALs [15]. Whereas a direct effect of neuronal activity in the CX and LALs on thoracic steering reflexes has been shown in cockroaches [16] [17] [18], the anatomical identity of the involved neurons and their place in the CX circuit is not clear. Additionally, how descending neurons encode motor commands on a population level is currently not well understood, but multiple recent studies have been able to dissect single neural circuits that underlie specific behaviours (e.g. [19] [20] [21]). One such behaviour, which has been examined in detail, is the pheromone-following behaviour of silkworm moths. Male silkworm moths display a highly stereotyped behavioural sequence when following a female’s pheromone plume. Upon first contact with the plume, the moth responds with a ‘surge’, that is a straight movement towards the source of the odour. When the odour plume is lost, ‘casting’ is initiated, during which the moths walks in a zig-zag pattern until it finds a new odour pocket (review: [22]). Early studies have identified several descending neuron types whose activity correlates with turning behaviour when a male moth orients in a pheromone plume [23]. Among these, the most notable are “flip-flop” neurons, which are bistable neurons that switch between a high-activity and a low-activity state in response to a trigger stimulus [24] [25] [23]. That is, the same stimulus can cause the neuron to increase or decrease its firing, depending on whether it is in the low or high activity state, respectively, when that stimulus occurs.

These neurons have post-synaptic terminals in the LALs, and their axons descend through the ventral nerve cord and synapse onto neck motor neurons, which in turn activate neck muscles that control head movements [26] [27]. Thus, if the left-descending flip-flop neuron is in its high-activity state, the left neck motor neuron and the left neck muscle are also active, causing the head and consequently the moth to turn left. Although this network has been described in the context of pheromone following, other studies have shown that flip-flop neurons can also be triggered by light flashes or sounds [24] [28]. It therefore seems likely that flip-flop neuron mediated steering may constitute a general form of targeted steering, independent of the stimulus modality that drives the behaviour [29].

In this study, we aim at evaluating whether a basic flip-flop neuron network can produce naturalistic steering in (a) a simulated odour plume and (b) when presented with visual input via a neural model of the central complex. To this end, we present a rate-coded and a spiking computational model of a simple flip-flop network. Both models follow the same connectivity patterns, but vary in their level of biological detail for the neurons. The rate model uses continuous-valued sigmoid neurons, while the spiking model uses leaky integrate-and-fire (LIF) spiking neurons. These are the two most common types of rate and spiking neural models, respectively. We examine these two models in order to determine in what ways the neuronal details matter. That is, the similarities in behaviour between the two models indicate aspects of the model where these implementational differences do not result in behavioural changes. As we will show, both models are effective at navigating towards a simulated pheromone source, so the low-level neural details do not matter for obtaining the essential functionality. However, we will also show that finer details of the behaviour resulting from the two models does differ, with the spiking model producing more realistic trajectories that we were unable to generate using the rate model. Importantly, the spiking model produces this more realistic behaviour while still following the same connectivity patterns as the rate model.

Of course, these two neuron models (rate-code and leaky integrate-and-fire) are not the only options for neuron models. More neural details can always be added, bringing the model closer to the real system. However, there are many potential details to add, and no consensus on which features are the most important ones to include. Indeed, part of the purpose of creating models using different types of neurons is exactly to find out which details matter.

This is why we chose to create our spiking neural model using the Neural Engineering Framework (NEF; [30]), which is a generic method for taking any neuron model and connecting the neurons such that they perform a particular algorithm. Instead of having a single rate-code neuron that can only produce an output that is a sigmoid function of the sum of its inputs, we can instead have multiple neurons of any type in a small group. By weighting the outputs of these neurons in different ways, the system can approximate a wider variety of functions, as opposed to just being able to compute sigmoids. This allows us to specify what computations we would like to be performed between neural groups, and the NEF will determine the precise weighting of each connection between those groups of neurons that will best approximate the desired computation.

Despite their differences, we show that both models reliably reproduce steering behaviour. Comparing the behaviour of our models to behavioural data from male silkworm moths, we find that a simple flip-flop based neural circuit is sufficient to replicate the basic characteristics of the moths’ paths. Furthermore, we demonstrate for the first time that the flip-flop network can work as a general steering network when combined with a computational model of the CX [14]. This study is therefore a step forward to close the gap between higher processing centres in the brain that make navigational decisions, such as the CX, and the thoracic motor circuits that ultimately move the insect.

## Materials and methods

To implement the spiking model, we used the software toolkit Nengo [33] which includes a software implementation of the NEF. All source code for both models is available at https://github.com/stanleyheinze/insect_steering.

### Models

#### Rate model

For the rate model, we use sigmoid neurons, with one addition described below for the “flip-flop” neurons. If the total input to the neuron is *J*, then the output *r* from the neuron will be

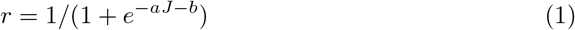

where *a* and *b* are the gain and bias constants for the neuron, respectively. The total input to the neuron is the weighted sum of the rates of the neurons connecting into that neuron

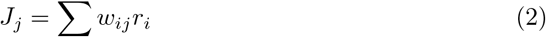

The weights *w* are either 0 (no connection), 1 (excitation), or −1 (inhibition), with gaussian noise of standard deviation 0.01 added when the model is created. We also add random noise 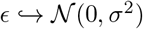 with *σ*^2^ = 0.02 to the input to each neuron on every time step. These two sources of randomness are meant to give some individual variation to the models. For the PBN and flip-flop neurons, we also add the neuron’s previous output to its own input, in order to allow for the sustained activity that has been described for these neurons. This gives them the following equation, where *r_t_* is the output for time step *t*.

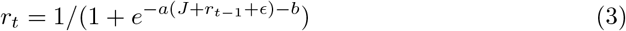

The neurons modelled here were physiologically and anatomically described in [24] [34] [23] [25]. We infer input and output regions of neurons from their anatomical appearance, i.e. smooth terminals are assumed to be inputs, while varicose terminals are assumed to be outputs. Two neurons are assumed to be connected if the input region of one neuron overlaps with the second neuron’s output region. The model connections are furthermore based on the network proposed in [23] with some small modifications.

The network consists of two pairs of neurons: A flip-flop neuron (FF) and a protocerebral bilateral neuron (PBN; Fig. 2A). Both cell types receive input either directly from the plume, or from the output neurons of the central complex (CPU1 neurons), when connected to the path integration network (PI). PBN neurons were proposed to provide bilateral inhibition between the two LALs [23] [35] and are therefore modelled to inhibit the contralateral FF neuron. FF neurons have excitatory connections directly onto the contralateral motor (Fig. 2B), based on the finding that FF neuron activity correlates with neck motor neuron activity [26]. For the FF neurons, we needed to add some mechanism to produce the flip-flop behaviour, where an input stimulus will switch a neuron from a high state to a low state. Since sigmoid neurons by themselves are too simple to produce this behaviour, we added a feature to the rate-code model where if the current output r is large (> 0.8) and the input is large (> 0.5), then the output of this FF neuron is reduced by 0.5 and the opposite FF is increased by 0.5. This produces the required flip-flop behaviour (Fig. 2B), but does not postulate a plausible mechanism whereby this behaviour is produced. We present a potential mechanism in the next section on the spiking neuron model.

**Fig 2.**
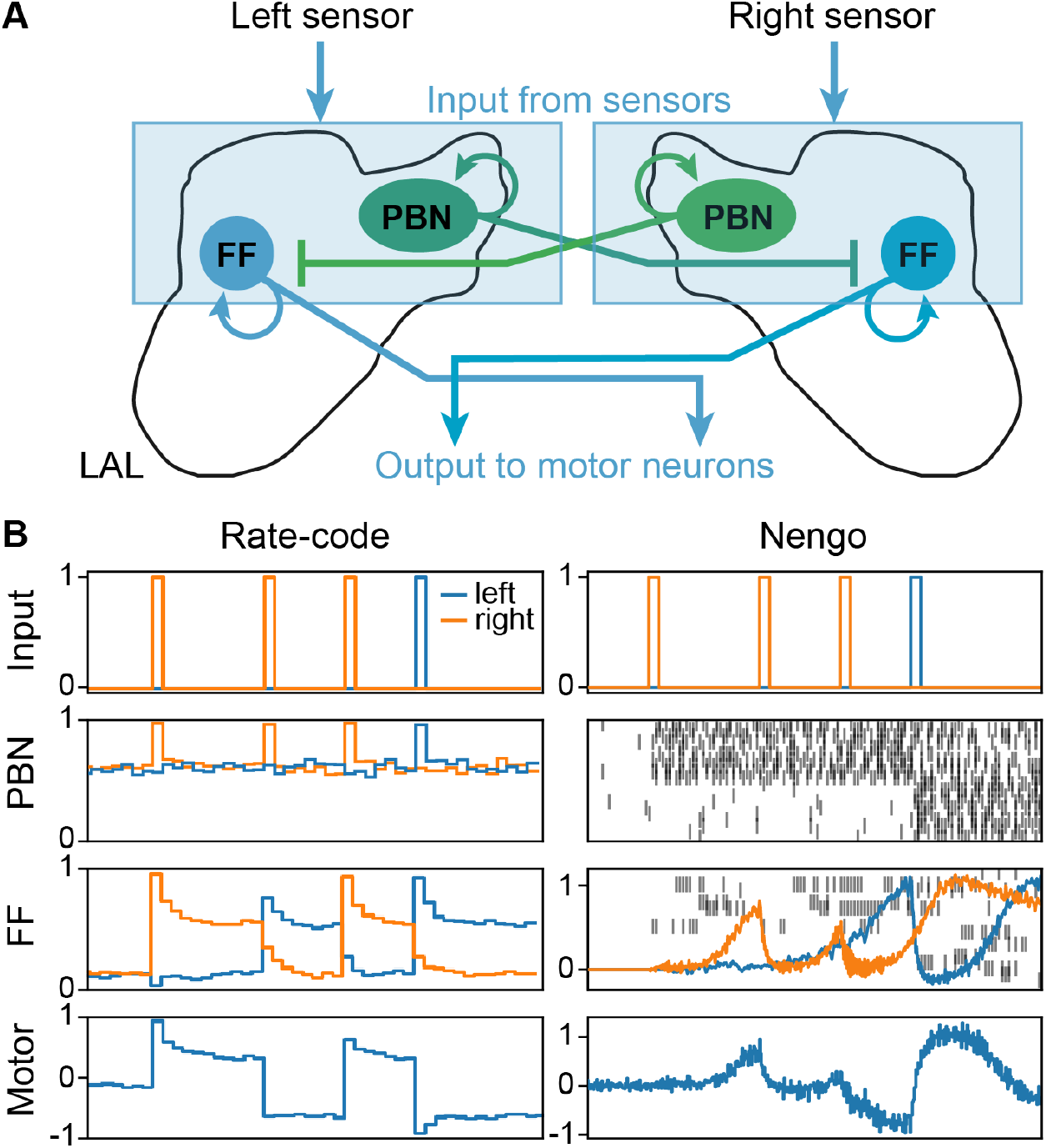
The flip-flop network. A: Schematic of the network. Protocerebral bilateral neurons (PBN) and flip-flop neurons (FF) both get input directly from the ipsilateral sensor. PBNs inhibit the contralateral FF. FF neurons activate the motor. LAL = lateral accessory lobe. Note that in the rate model, a state switch in one FF neuron explicitly affects the other neuron, which is not reflected in the anatomical connections shown. B: Rate-code and Nengo network responses to the same artificial input. For the Nengo PBN and FF neurons, the vertical lines show the activity of a random subset of the internal components that are used to construct the neuron model, with the components of the right neuron at the top and the left neuron at the bottom. The output from the FF neurons is shown as the orange and blue lines, and is computed as the weighted average of the internal activity. The rate-code model has an explicit rule that when one FF is forced to the down state (at the second input pulse), the other is forced into its up state. The internal components of the Nengo FF neurons cannot approximate this part of the flip-flop behaviour, but even without this it still settles into an asynchronous activity pattern (after the third input). For longer timeframe, see S1 Fig.

#### Nengo model

Our second model is intended to explore how the basic operations of the rate-code neural model described above - which uses sigmoid functions and flip-flop logic - can be constructed out of more biologically realistic components. Specifically, we use the Neural Engineering Framework (NEF; [30]) to define sets of spiking neurons with weighted connections that can approximate the desired functions. Note, the spiking model presented here is not meant to be an accurate model of the lateral accessory lobes; rather, it is just meant to be more realistic in its basic operations than the rate-code model, and to allow us to see what changes in behaviour result from this change in implementation.

We use groups of 100 leaky-integrate-and-fire neurons to represent each of the single neural units used in the rate-code model. We use least-squares minimization to find the optimal connection weights that will make the group of spiking neurons behave as closely as possible to their rate-model equivalents. The same process is used to approximate both the sigmoid input-output function and the flip-flop behaviour. In the latter case, we assume recurrent connections between the neurons in the group representing the flip-flop and find the appropriate weights between them to give the desired property that an input will switch the output between high and low firing. Note, however, that we are not suggesting the flip-flop neurons must have recurrent connections, but rather that these neurons have a sufficiently complex internal state that such feedback is necessary to capture their properties.

We call the model a Nengo model because we used the Python software package Nengo (Bekolay et al., 2014) to automate the process of building our model. All source code for our model is available at https://github.com/stanleyheinze/insect_steering. The resulting approximation is not perfect, and as can be seen in the right-hand column of figure 2, and produces a notably different time course of response compared to the rate model, although the key qualitative characteristics are maintained.

One crucial difference between the rate-code and Nengo models is in how synchronization between the two flip-flop cells is achieved. In the rate-code case, we have explicitly added a rule that forces the contralateral flip-flip to its up-state when an input pushes the ipsilateral flip-flop to the down-state. However, the purpose of the Nengo model is to build up the desired behaviour out of more basic components. The components inside one flip-flop neuron can successfully switch the state of the flip-flop in the Nengo model. However, there is no way for the basic components inside one flip-flop neuron to affect the contralateral neuron. This means that the Nengo model has mostly the same behaviour as the rate-code model, with one exception: when one flip-flop changes to its down-state, it does not force the other flip-flop into its up-state. The asynchronicity between the two flip-flop neurons is therefore forced in the rate-code case, but emerges in the Nengo case.

### Experimental situation 1: Following an odour plume

Our model is directly inspired by the flip-flop neurons that have been implicated in pheromone tracking in moths, hence we first evaluate its ability to control the behaviour of a simulated agent in an odour plume (figure 3A). The plume was simulated by the python-based module pompy (https://github.com/InsectRobotics/pompy, by Matthew Graham, Insect Robotics Group, Edinburgh University), using parameters that are appropriate for a gypsy moth pheromone plume ([36]; see table 1 for plume parameters). The plume was dispersed by a weak constant wind flowing from the direction of the odour plume source towards the agent’s starting point. The agent was equipped with two fronto-lateral sensors which were designed to mimic antennae. The antenna size and angle (away from the centre line) from each other was adjusted to match real moths ( [37]; see table 1 for moth parameters). We set a maximum antenna sensitivity, above which the response is saturated, i.e. any higher concentration does not elicit a stronger response. We then choose a maximum response value to correspond to this level of concentration, and scale the response linearly between this value and zero for lower concentrations. These two parameters were adjusted by hand. At each point in time, the network processes the input and makes a steering decision, which is applied to the centre of mass of the agent and scaled by the maximum rotation speed (table 1), analogous to [14]. The agent’s walking speed was determined by an acceleration and a drag parameter, analogous to [14] (see table 1). The agent’s acceleration setting in the simulation was adjusted to match the actual walking speed of moths [37]. The resulting forward speed was not influenced by the headwind, but agents were given a weak tendency to turn upwind in the absence of plume input, in accordance with observations in silkworm moths [22]. Unless otherwise specified, all parameters were tuned manually to match the behaviour of real moths as closely as possible. Thus, certain arbitrary parameters of the models, such as the maximum output value for that antenna and the maximum rotation speed were set such that the resulting tracks approximated the tracks of silkworm moths as reported in [38].

**Fig 3.**
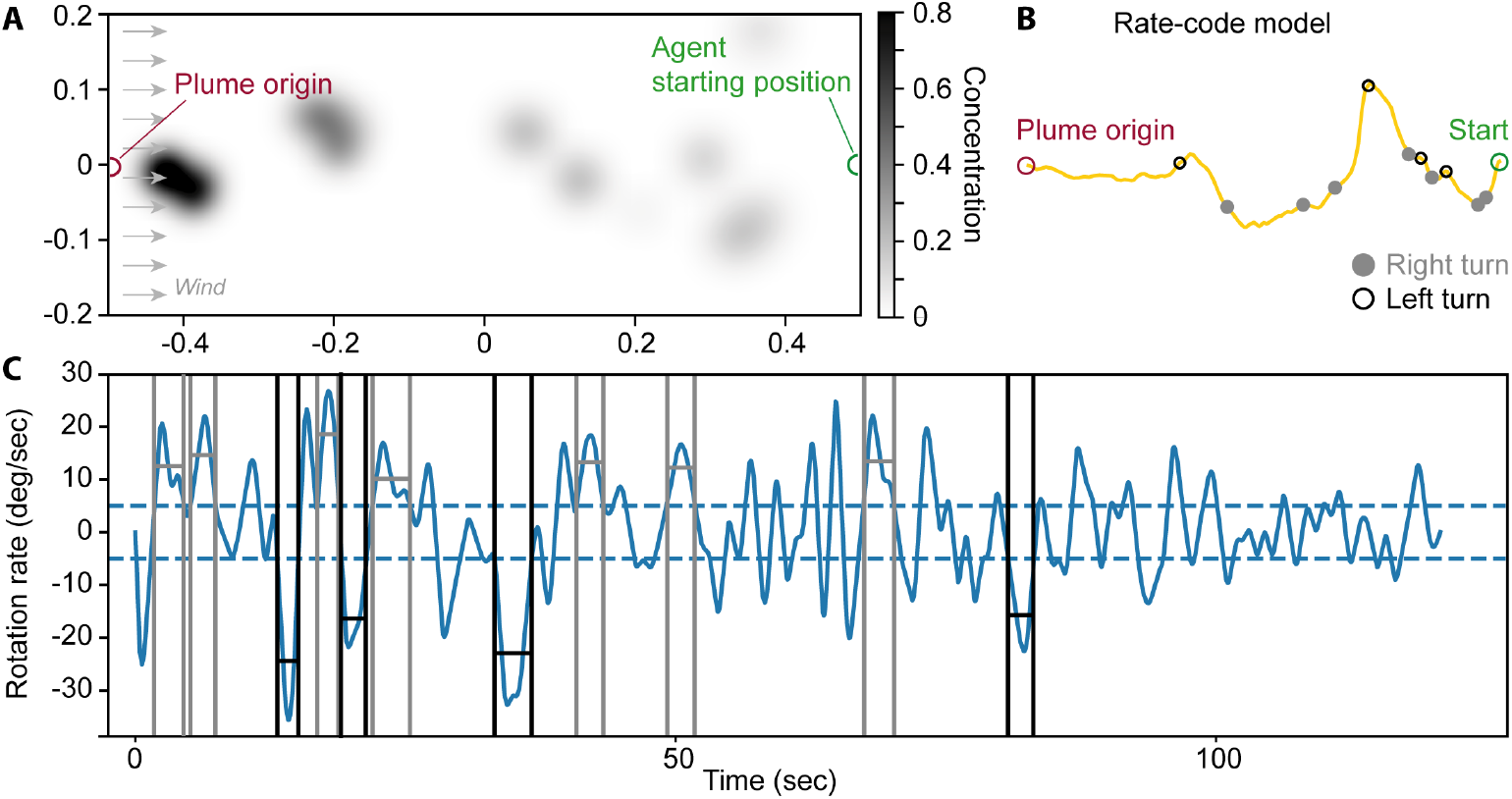
Overview over the odor plume experiment. A: The pheromone plume is released from the source and dispersed by wind. Shown is one frame of the plume, with the grey value reflecting the pheromone concentration on a scale from 0 to 1. The agent is expected to navigate towards the source using the odor plume. In the absence of input, the agent turns upwind. B: Typical path of the rate-code model, with right turns indicated by filled grey circles and left turns indicated by empty black circles. C: The rotation rate defines whether a rotation is classified as a turn. Detected turns are marked (right turn = grey, left turn = black). Definition as in [38].

**Table 1.** Simulation parameters.

| Parameter | Value |
| --- | --- |
| Number of LIF components to implement one neuron | 100 |
| Random noise added to neurons | $\mathcal{N}(0, 0.02)$ |
| Wind speed | 2 m/s |
| Plume puff release rate | 50 Hz |
| Plume puff initial radius | 0.1 m |
| Antenna size | 7.5 mm |
| Antenna angle | 45° |
| Antenna maximum sensitivity | 100 |
| Antenna maximum output value | 3.0 |
| Moth maximum rotation speed | 8.0 rad/s |
| Moth acceleration | $0.2m/s^2$ |
| Moth drag | 0.5 m/s |
| Moth turn-into-wind rate | 0.1 rad/s |

Ando et al. (2013) [38] presented a robot that was steered through an odor plume by an on-board moth walking on a track ball. The moth’s movement on the trackball was translated into wheel speeds for the robot. The authors presented trajectories for both the moth-controlled robot and the moth only in the odor plume. Here, we adjusted the parameters of this model such that the agent’s trajectories were similar to the moths’ trajectories presented in [38], based not only qualitatively on the trajectories, but also quantitatively on the turn duration, turn angle and turn velocity.

Using their robot, Ando and colleagues performed further experiments where they added a bias to the turning of the robot. That is, a constant signal was added to the left (or right) wheel, while the moth was controlling the robot. This causes the moth to drift to the edge of the pheromone plume, but they show the moth is able to compensate for this and continue to follow the plume. We performed the same experiment in our simulation by adding a constant bias (in rad/s) to the moth rotation, causing the simulated agent to have an extra tendency to turn in a particular direction. Ando and colleagues show that moths plume-following behaviour is robust to this sort of manipulation, and that the resulting paths tend to follow along the edge of the plume.

Behavioural measurement thresholds were defined in accordance with [38], to allow for comparing the models to data from male silkworm moths. A turn was identified if the agent’s turn duration was larger than 0.5 s, the turn angular velocity was larger than 5 deg/s, and the turn angle was larger than 30° (figure 3B, C; [38]). Loops were detected based on their high rotation rate, above 30 deg/s for at least 5 s. An experiment was considered successful if the agent arrived within 5 cm of the goal. All data was analysed in Python 3.5.5.

### Experimental situation 2: Path integration

A second motivation for our model was to understand how output from the central complex is translated to steering behaviour. Specifically, we connected our steering system to a previously developed CX path integrator model using sigmoid neurons [14]. This model is based on the neuroanatomy of the central complex, and we take the output neurons from the model (CPU1; figure 6A) and project them to the lateral accessory lobes in our model, where they may interact with the flip-flop descending neurons as well as the protocerebral bilateral neurons modelled here. Since there are 8 CPU1 cells per hemisphere whose summed activity is thought to activate the motor, but only one flip-flop neuron in our model, the activity of all 8 neurons was summed by projecting onto the same flip-flop cell.

In the plume experiments, the value of each sensor was between 0 and 1, so that the difference between the two sensors could also fall between 0 and 1. However, the output from the path integrator had a narrower spread (0.5 to 1). While our models worked with that smaller difference, we also re-scaled the path integrator output to a scale of 0 to 1 to test whether this would improve the models’ behaviours.

To evaluate the behaviour in this situation, we used exactly the same simulator as before (pompy), but removed the odour plume. We then caused the agent to take a random exploratory path by setting its rotation rate to be a Gaussian white noise process with *σ* = 0.1rad/s while moving forward at a constant speed (see table 1), for 15 seconds in total. The agent then attempted to return directly home using the CX output combined with our steering model. The simulation was then continued for another 40 seconds. An experiment was considered successful if the agent arrived within 5 cm of the starting location.

## Results

In order to determine how realistic the behaviour of our model is, we compared the simulated tracks qualitatively and quantitatively to data from silkworm moths (*Bombyx mori*, originally published in [38], Fig. 4). The total turn duration of the rate-code model falls within the standard deviation of real moth data but was slightly higher for the nengo model (Fig. 4A). The total turn angle of both models fell within the standard deviation of real moth data (Fig. 4C), while the mean turn velocity of both models was 10-15 % lower compared to real moths (Fig. 4B). However, both models performed well with respect to finding the origin of the plume, with a success probability of 0.84-1.0 (Fig. 4D). Qualitatively, the tracks of both models display clear zigzagging (Fig. 4E, F), with the rate-code model having overall straighter paths than the nengo model. Looping behaviour is seen in the nengo model and, to a lesser extent, in the rate-code model (Fig. 4I). Since moths have been described to perform loops when they lose the odour plume and search for a new pocket of odour, we checked the sensor values during looping. Loops were detected automatically based on their high rotation rates, allowing us to divide the trajectory into looping and non-looping stretches (Fig. 4H). Analysing the difference in value between the two sensors during looping, we find that looping occurs proportionally more often when the two sensors have similar or equal values (Fig. 4G), suggesting that looping emerges from the model when the sensor data is non-directional, which is consistent with real moth behaviour. Overall, when comparing the rate-code and nengo models to moths, we find that both models replicate real moth data reasonably well.

**Fig 4.**
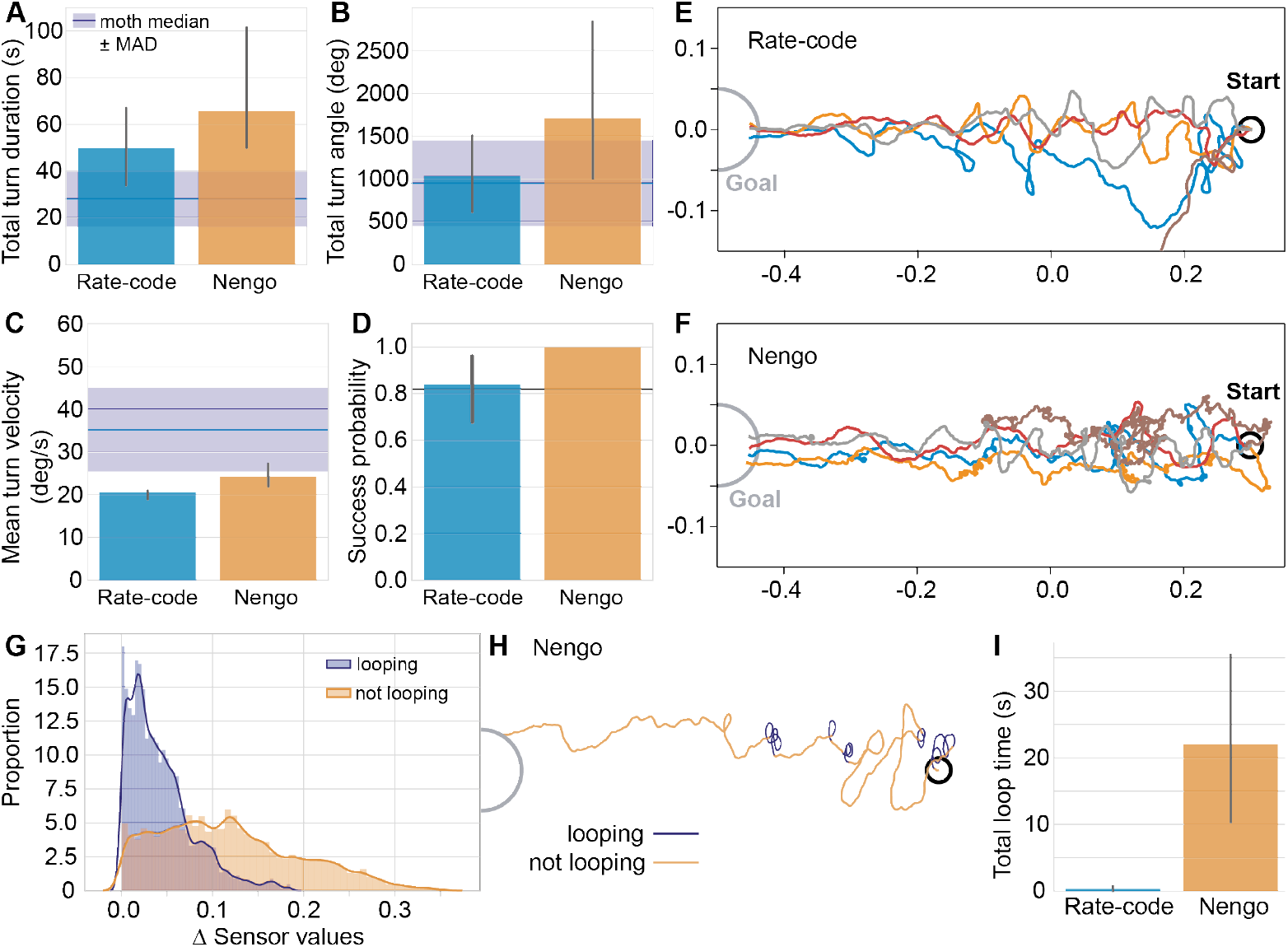
Behavior of both models in a simulated odor plume. A-D: Comparison of the rate-code and Nengo model to data from silkworm moths. Bar plots represent the median, with the error bar giving the bootstrapped 95 % confidence interval. The moth mean ± standard deviation (black line and grey area) and the moth median ± median absolute deviation (dark blue line and blue area) are given for comparison. Moth data reproduced with permission from [38]. E-F: Example trajectories for the rate-code (E) and Nengo (F) models. G: Difference in sensor values during looping. During loops, the values of the two sensors are very similar or equal. H: Example trajectory of the Nengo model displaying automatically detected loops in dark blue, and non-looping stretches in orange. G. The Nengo model displays substantially more looping than the rate-code model. For additional paths, see S2 Fig

A further way of testing how well our models replicate real moth behaviour was to add a turn bias to the simulation. When given a turn bias, silkworm moths were shown to track the edges of the odour plume instead of the centre [38]. This was also the case for our models (Fig. 5). When analysing the angle between the current position of the agent and the source of the odour plume, we found that the models shift away from the centre of the plume already at a turn bias of 1 rad/s (Fig. 5A, B). With increasing turn bias, the models’ success rate decreases, and at a turn bias of 3 rad/s, no agent simulated by the rate-code model reaches the goal. The nengo model is more robust but starts failing at a turn bias of 5 rad/s (data not shown). While these results are more difficult to compare quantitatively to real moth data owing to the different ways of implementing the turn bias, it is clear that qualitatively the models behave very similarly to real moths, as they track the edge of the plume rather than the centre when given a turn bias.

Having established that the simple flip-flop networks are able to reliably replicate several characteristics of male silkworm moth behaviour, we then proceeded to test whether the models could also take CX output, rather than odour plume information, as input signals. For this experiment, we used a computational model of the CX that computes path integration in an anatomically constrained network (Fig. 6A; [14]). In short, path integration is a computation that combines, at each time step, the current heading of the animal (represented in TB1 neurons, Fig. 6A) with its forward speed (represented in TN neurons, Fig. 6A). The resulting vector (updated in the CPU4 memory loop, Fig. 6A) points in the direction of the path’s origin, and the length of the vector represents the distance of the animal from that origin. Thus, an animal that continuously updates this vector during an outbound path has the possibility to return back to its origin in a straight line. Once the animal decides to return, the desired heading (encoded in the CPU4 vector) is compared to the current heading (TB1 neurons), and mismatches between the two are transferred to an unspecified motor via CPU1 neurons. As CPU1 neurons project from the CX to the LAL, it is plausible that they interact with flip-flop neurons there. Importantly, steering signals are encoded as an imbalance between the summed activity of all CPU1 neurons in the right hemisphere and those in the left hemisphere. We therefore used the summed CPU1 activity as input to the flip-flop network. Using the same model parameters as for odour-plume experiments, both the rate-code and the nengo model steer an agent back to its origin based on path integrator output (Fig. 6B), albeit with a lower success rate than the ideal path integrator (Fig. 6D). We assessed the accuracy of homing by analysing the orientation of the agent relative to the origin, where 0° means that the agent is perfectly tracking along the straight line between the end of the outbound path and its origin (Fig. 6C). Without any connections to the steering model, the ideal path integrator peaks at an orientation of 0° and has a standard deviation of 15.4°. The nengo model also peaks at 0° but has a wider standard deviation of 67.9°. Interestingly, the rate-code model has one peak at 0°, a standard deviation of 50.2°, and two additional peaks at ± 90°.

Due to this odd distribution of orientations, and due to the relatively low success rate for the two models (0.7 and 0.57, respectively), we examined different ways of connecting the path integrator to our steering system. After all, there is no a priori reason for expecting an output value of 0.5 from the path integrator to mean the same thing as a 0.5 from the odour detection system. However, we do not want to postulate complex neural mechanisms between these two systems. Two simple things to adjust would be the gain of this connection (which would correspond to increasing the number of synapses, or moving the synapse closer to the spike initiation zone) and adjusting the bias current (which would correspond to changing the threshold at which the neuron will fire). While neither of these on their own significantly improved behaviour, we found that adjusting this gain and bias such that a path integrator value of 0.5 is mapped to a 0 input to the steering system and a value of 1.0 stays at 1.0 (and intermediate values are linearly interpolated between these) greatly improved performance while keeping the same qualitative effects (Fig. 6). When re-scaling the path integrator output to a scale between 0 and 1, we find that both models are significantly more successful as well as more accurate in tracking along a straight line back to the origin (Fig. 6E, F, H). At a sensor difference of 0.5 or above, the rate-code model’s percentage of successful runs increases from 0.7 to 0.78, and the Nengo model increases from 0.57 to 0.92 (Fig. 6H).

## Discussion

We modelled a simple flip-flop network based on neurons that have been described in detail in the silkworm moth. The computational model has only two pairs of neurons: the flip-flops, which are bistable neurons, and the PBNs, which provide inhibition between the two hemispheres. This network was modelled both as a rate-code model and as a spiking nengo model. Surprisingly, both models were able to replicate the behaviour of real moths in an odour plume reliably, despite their simplicity. We also tested whether this simple steering network could serve as an interface between the CX and downstream motor centres by combining it with the CX path integrator network [14]. We could show that both the rate-code and the nengo model can take input from the path integrator and use it for steering towards a target, and that the efficiency of steering depends on the scaling of the input into the system. In the following, we will discuss our findings with respect to the models, their behaviours and the predictions and conclusions we can draw from these experiments specifically as well as computational models of insect neural networks in general.

### Rate vs. Nengo model

When comparing the rate-code and the Nengo models, both produced very similar overall plume-following behaviours, despite the difference in complexity. This shows that rate-code models are, to a certain degree, a valid way of modelling neural networks, despite the many simplifications they involve by default. However, the more subtle behaviours of the system seem to be more realistic in the Nengo version, in particular for looping behaviour and for the influence of turning bias.

For the looping behaviour, Figure 4 E and F show that the Nengo model provides frequent tight looping movements that are much less common in the rate model. We were unable to find ways of modifying the rate model to create these tight loops while still being able to successfully follow the plume. Similarly, for the behaviour when a turning bias is introduced (Figure 5B), the Nengo model produces the same edge-following behaviour observed in the original experiment, while the rate version does exhibit some shift, but with a wider spread. Overall, this indicates that the Nengo model produces more realistic behaviour, with its approach of using low-level components to approximate more complex neural functions. For this reason, we believe the Nengo model is worthy of future study.

**Fig 5.**
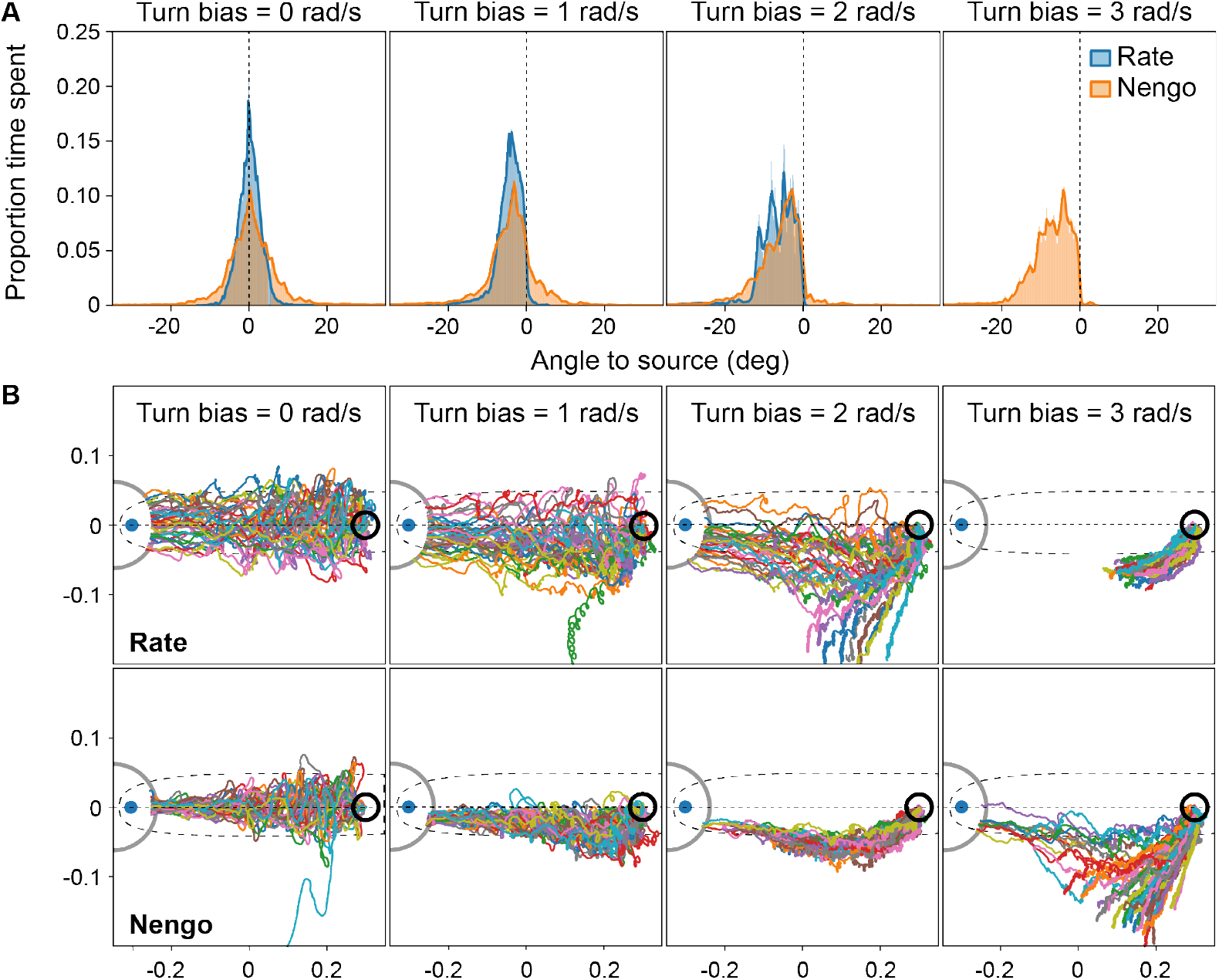
Turn bias. A: Proportion of time the agent spent at a specific angle relative to the source of the odor plume. Only successful trials were included. When a turn bias was added to the left motor, the agents shifted away from the center of the plume towards the plume’s edge. At a turn bias of 3, the rate-code model failed consistently. B: Paths of the rate-code and Nengo models with increasing turn bias. N = 40 per condition. Dotted line = center of the odor plume; dashed outline = area of odour concentration of at least 10 percent of the maximum detectable level, averaged across 3000 timesteps; blue dot = plume origin; grey circle = area around plume origin that needs to be reached in order to count as success.

**Fig 6.**
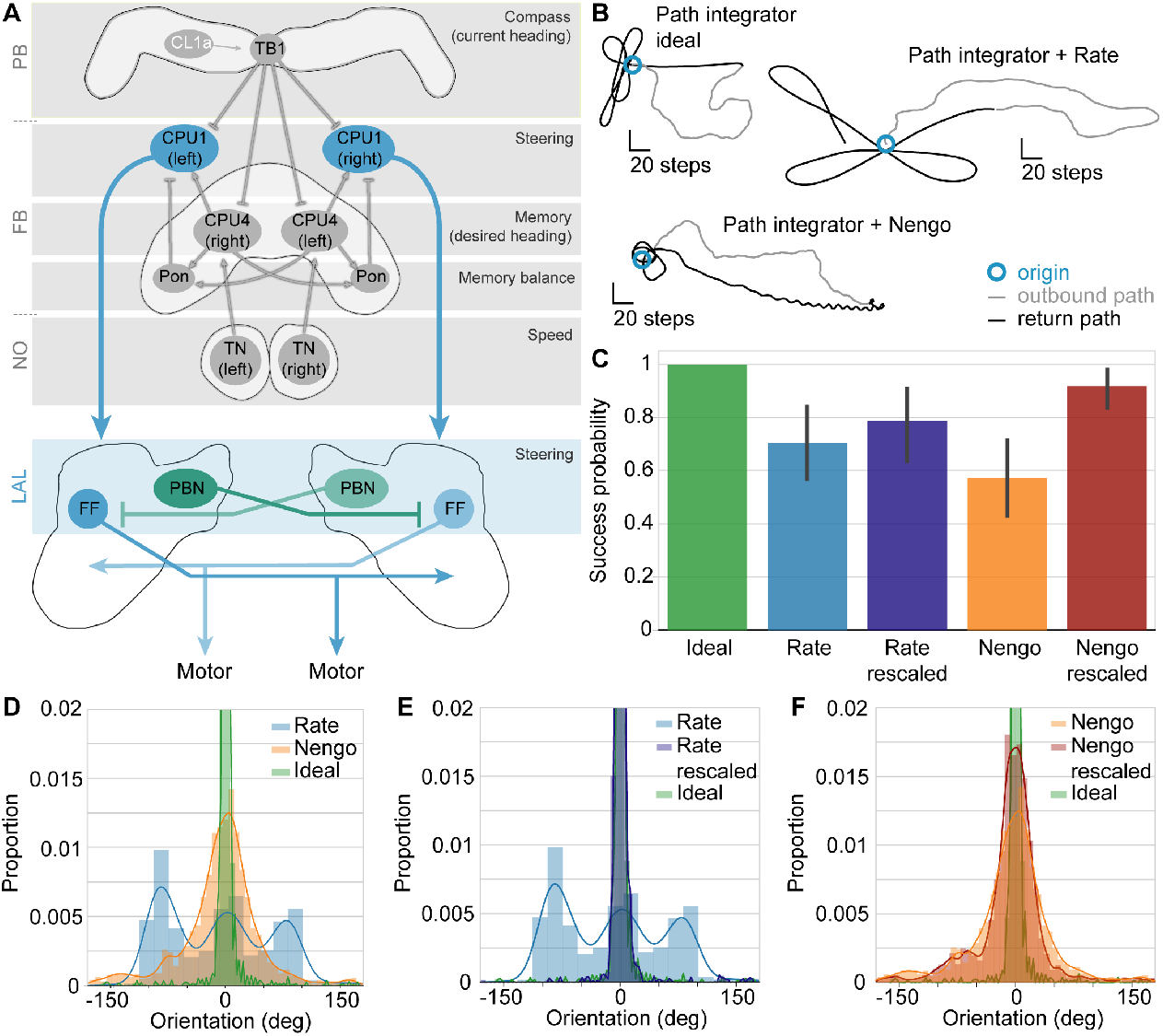
Integrating the flip-flop models with the central complex path integrator. A: Schematic of the integrated network. The path integrator receives compass input via CL1a neurons, and speed input via TN neurons. TB1 neurons compute the current heading direction. CPU4 and pontine (Pon) neurons form a memory loop which integrates the current heading direction with the current speed, resulting in a vector that points to the path’s origin. When the agent wants to return home, CPU1 neurons compare the current heading (represented in TB1 neurons) to the desired heading given by the CPU4 home vector. CPU1 neuron output is then fed directly into the flip-flop networks, with the same input driving both the inhibitory protocerebral bilateral neurons (PBN) and the flip-flop neurons (FF). B: Example paths of the path integrator without a steering network (ideal), with the rate-code model, and with the Nengo model. C: Success rate of the integrated models compared to the ideal path integrator. The success probability of both models increases when the CPU1 output is rescaled to a range of 0 to 1. Error bars represent the bootstrapped 95 % confidence interval around the median. D: Proportion of time the models spend at a certain angle relative to the path’s origin. E: With rescaled input, the distribution of the rate-code model becomes almost identical to that of the ideal path integrator. F: With rescaled input, the distribution of the Nengo model has a smaller spread around 0° and becomes more similar to the ideal path integrator. N = 50 for all path integrator experiments. Only successful runs were considered in D-F.

### Steering network

The simple steering network presented in this paper appears to be sufficient to mediate efficient and realistic steering behavior. However, many more neuron types than the two modelled here were described to be part of the pheromone-responsive network in the LALs of the silkworm moth [34] [25] [23] [39] [40]. Thus, while producing realistic steering behavior, our models are a gross simplification of the actual neural circuit that likely computes pheromone-following. For example, two neuron types that were proposed to play a role in steering are GII-A and GII-C descending neurons [23] [35]. We found that they were not necessary to produce the behavior presented in this paper (S2 Fig). Including more neuron types in this steering network was beyond the scope of this paper, but will certainly refine the resulting behavior and potentially lead to unexpected emerging behaviors (see discussion of looping behavior below).

A major open question is how flip-flopping can be achieved in a real neuron. In the Nengo model presented here, we use an optimization method to combine low-level components, which consist of voltage build-up and spikes, to approximate the flip-flop behavior. While this produces more realistic behavior than the idealized rate-based approach, more realistic details could be added. In particular, various models of neural bistability currently exist (e.g. [41] [42]) that might serve as the basis for a more accurate model of these flip-flop neurons. Importantly, since these theories postulate internal mechanisms as the basis of the flip-flop behavior, we can use the same Nengo software and the same function approximation process to generate these more accurate models, and see if the resulting behavior is also more accurate.

### Model behavior

When evaluating the behavior of both models, the turn angle, turn angular velocity and turn duration agree well with values reported for male silkworm moths and fall within the mean ± one standard deviation of moths [38]. The high positive deviation of the Nengo model’s turn duration (Figure 4A) can be explained by this model’s tendency to loop, leading to more and longer turns. The rate-code model also produced looping behavior, but at a much lower rate than the Nengo model. Interestingly, loops occurred when the difference between the two sensors’ values was very low, up to 0.1. While many insects perform loops (e.g. [43] [44] [45]), this locomotion strategy would not be expected in simple taxis behaviors, in which the animal navigates purposefully towards or away from a sensory cue. In this situation, a very small difference between the two sensor values would be expected to elicit a straight walk, with the aim of keeping the difference as small as possible. On the other hand, looping allows the insect to sample its entire immediate environment, which may be especially useful if the insect orients relative to an external cue, or towards the source of an intermittent cue such as an odor plume. For example, fruit moths perform a loop as a search mechanism when they lose the odor plume [43]. Dung beetles perform a circular dance on their dung ball to sample their environment and to take a ‘snapshot’ of skylight cues before rolling their dung ball in a straight line relative to these cues [6] [44], and perform another dance after losing their bearing. Desert ants use a similar strategy to learn their visual surroundings during learning walks [45]. Looping therefore appears to be a robust strategy to sample and learn the immediate sensory environment, as well as to re-acquire a sensory signal that has been lost. It is intriguing that the network architecture of our simple model gives rise to looping as a search behavior. This finding suggests that looping may not be an active search strategy, but one that simply emerges from the connections within the network in situations where sensory information is ambiguous. Exactly how this behavior emerges is outside the scope of this paper, but our initial hypothesis is that it is a side effect of the timing behavior of the Nengo model. The Nengo model consists of LIF neurons and synapses, organized to approximate flip-flop neurons and sigmoid neurons. But these LIF neurons and synapses have intrinsic timing (membrane time constants and post-synaptic time constants, respectively). These are both on the order of tens of milliseconds, allowing the neural system to hold information over time. This may allow the moth to continue its tight loop during times when the odor plume is intermittent in a way that the rate model (with its lack of temporal dynamics) does not.

### Central complex output can be used for steering

In addition to steering towards the source of an odor plume, we could show that the models also steer well when getting input from the CX path integrator (PI) network published in Stone et al. (2017) [14]. The PI network itself computes a home vector based on visual information, using a visual compass signal for estimating its current heading, combined with optic flow to estimate its current speed. Both types of information are integrated in a memory loop and result in a vector that always points to the agent’s point of origin. When the agent wants to return home, it can use the vector to return there in a straight line by comparing its desired heading to its current heading and adjusting for any mismatches. Thus, the output from this network is in essence a steering signal that represents the difference between the intended heading and the current heading. The output is asymmetric between the right and left hemisphere, depending on whether the agent needs to correct to the right or to the left.

Our steering models can take this input and steer the agent towards its point of origin, using the same parameters that were used for odor-based steering. This was surprising, considering that the sensory input experienced in an odor plume is quite different from the input provided by the path integrator. Odor plume input is intermittent and varies at a high temporal frequency between 0 (no input at all) and 1 (the highest odor concentration possible), whereas path integrator output is constant, changes smoothly without sudden jumps, and ideally varies within a relatively small range around the intended heading. As the steering model was calibrated using odor plume input, it was not unexpected that the steering network steered less reliably using path integrator output, with the rate-code model in particular often steering at a 90° angle to the home vector. Re-scaling the difference to a range between 0 and 1 improved the success rate and accuracy of both models.

The finding that the steering networks can steer based on CX output shows that in principle, the flip-flop network is not restricted to using odor cues but can use signals from other modalities as well. One important limitation is that the input signals need to be directional, that is, there must be an imbalance between the signals in the right and left hemisphere. Here we specifically tested input signals that are derived from visual cues, but there is no reason that the input should be restricted to olfactory and visual cues alone. This agrees well with data from the silkworm moth, whose flip-flop neurons are known to switch state in response to odor cues as well as light flashes. Furthermore, bistable neurons of a similar morphology were found to switch state in response to auditory input in the cricket [28]. These data, taken together with our findings, make a strong case that the flip-flop network is not specific to mediating odor plume-following behaviors, but can take multimodal input, including input from the heading direction network in the CX, and produce robust steering. Our data therefore supports the idea that this neural network may work as a general purpose steering network at least in the context of targeted orientation behaviors (excluding taxis behaviors).

### Predictions

Our analysis generates several interesting and testable predictions. First and foremost, CPU1 neurons that project from the CX to the LAL are expected to have either direct or indirect excitatory synaptic connections with flip-flop neurons. To our knowledge, only one similar connection has been described so far: In the fruit fly, a CPU1 neuron (PF-L in *Drosophila* nomenclature) was shown to synapse onto bilateral LAL interneurons [46] (see also [47]. However, whether CX output neurons also project onto descending neurons in the LAL remains unknown. Finding the interaction sites between CX output neurons and LAL descending neurons, such as the flip-flop neurons modelled here, will be an important step towards understanding how the CX controls behavior.

Secondly, we propose that the flip-flop network does not only underlie olfactory steering, but that it can be a multimodal steering network. If this is correct across insects, we would expect flip-flop neurons to switch state in response to any stimulus that elicits targeted locomotion. Here, we will discuss three examples of targeted locomotion: straight line orientation, migration and path integration.

Dung beetles perform short-distance, straight-line orientation when rolling their ball away from the dung pile [48]. To keep their path straight, they rely on skylight cues such as the position of the Sun, the polarization pattern of the sky, and the sky’s spectral gradient [5] [49] [50] [51]. These cues are integrated in the CX to generate a current heading [44], which can be used to steer the animal along its straight path and adjust for deviations. We therefore expect that dung beetle flip-flop neurons should respond to a sudden change of the skylight cue, such as a sudden rotation of the polarization pattern, with a state change.

When it comes to long-distance migration, the Monarch butterfly and the Bogong moth are well-known insect models for diurnal and nocturnal migration, respectively [52] [53]. The Monarch butterfly uses a time-compensated Sun compass to migrate from its breeding grounds in North America to overwintering regions in central Mexico [7]. Additionally, it can use the geomagnetic field [54]. The Bogong moth uses the geomagnetic field in combination with visual landmarks to migrate from its breeding grounds in southern Queensland and western New South Wales (Australia) to its overwintering sites in the Australian Alps [8]. If the Monarch butterfly or the Bogong moth get off course and miss their target, they likely perish, thus precise course control is essential during their migration. The CX path integration network has been proposed to be a possible substrate for computing long-distance migration [10], and Sun compass information is also processed in the CX [55], thus making it likely that the resulting steering commands are passed on to LAL descending neurons. We would expect flip-flop neurons in the Monarch butterfly and the Bogong moth to switch state in response to sudden changes in the skylight cues or landmark configuration that they use to orient. Furthermore, many flying insects use optic flow for flight control, including moths [56] [57] [58] [59]. One might therefore expect flip-flop neurons to also respond to optic flow with a state change.

Finally, path integrating ants and bees are obvious targets for measuring flip-flop neuron responses, considering that we use the PI network as an input for our steering models. However, since the flip-flop neurons are not driven directly by sensory input that could be controlled in an experimental situation, but rather by a memory state, it is more difficult to test how flip-flop neurons respond during homing. One could however test optic flow cues and compass cues separately, to dissect how the different components of the path integrator drive the flip-flop neurons. Alternatively, it may be possible to perform extracellular tetrode recordings from flip-flop neurons during natural homing on a track ball [60].

## Conclusions and outlook

Of course, motor control mediated by descending neurons is much more complex than the simple model presented here. In the silkworm moth, several other neuron types were described to play a role in pheromone-mediated steering, which can be added to the model for increased complexity and biological relevance. Additionally, complex motor patterns are most likely mediated by not just one cell type, but by a population code across a number of descending neurons [15]. Creating larger and more integrated models of this form is a useful tool for a more complete understanding of these complex interactions.

We believe that the model we have developed and presented here is one small step towards understanding the connection between the heading direction system in the CX and downstream motor centers. Importantly, the approach we have taken to develop this model is flexible and suitable for a wide range of model features. While this is the most complex insect-based model developed using the Nengo neural modelling software, Nengo has also been used for a wide variety of mammal-based models, including Spaun, the first large-scale functional model of the human brain [61]. This involved modelling 30 different brain areas, including motor cortices, primary visual areas, the basal ganglia, and the thalamus, as well as modelling a simple environment for interaction. While modelling insect brains offers different challenges than mammalian brains, we believe our work has shown that this sort of large-scale model is possible, and can lead to more realistic behavior than some traditional modelling approaches.

Developing a more complete model cannot be done by a single group of researchers. We have made our model freely available at https://github.com/stanleyheinze/insect_steering, and we hope that a community of researchers can, over time, add additional neuron types and neural system to increase the complexity of the model and advance our understanding of this general steering system in insects.

## Acknowledgments

We would like to thank the organizers and participants of the Nengo summer school 2016, in particular Ben Morcos and Xuan Choo, for contributing to the initial moth model and for helpful discussions. We are also grateful to Noriyasu Ando for sharing his data with us.

## Supporting information

**S1 Fig.**
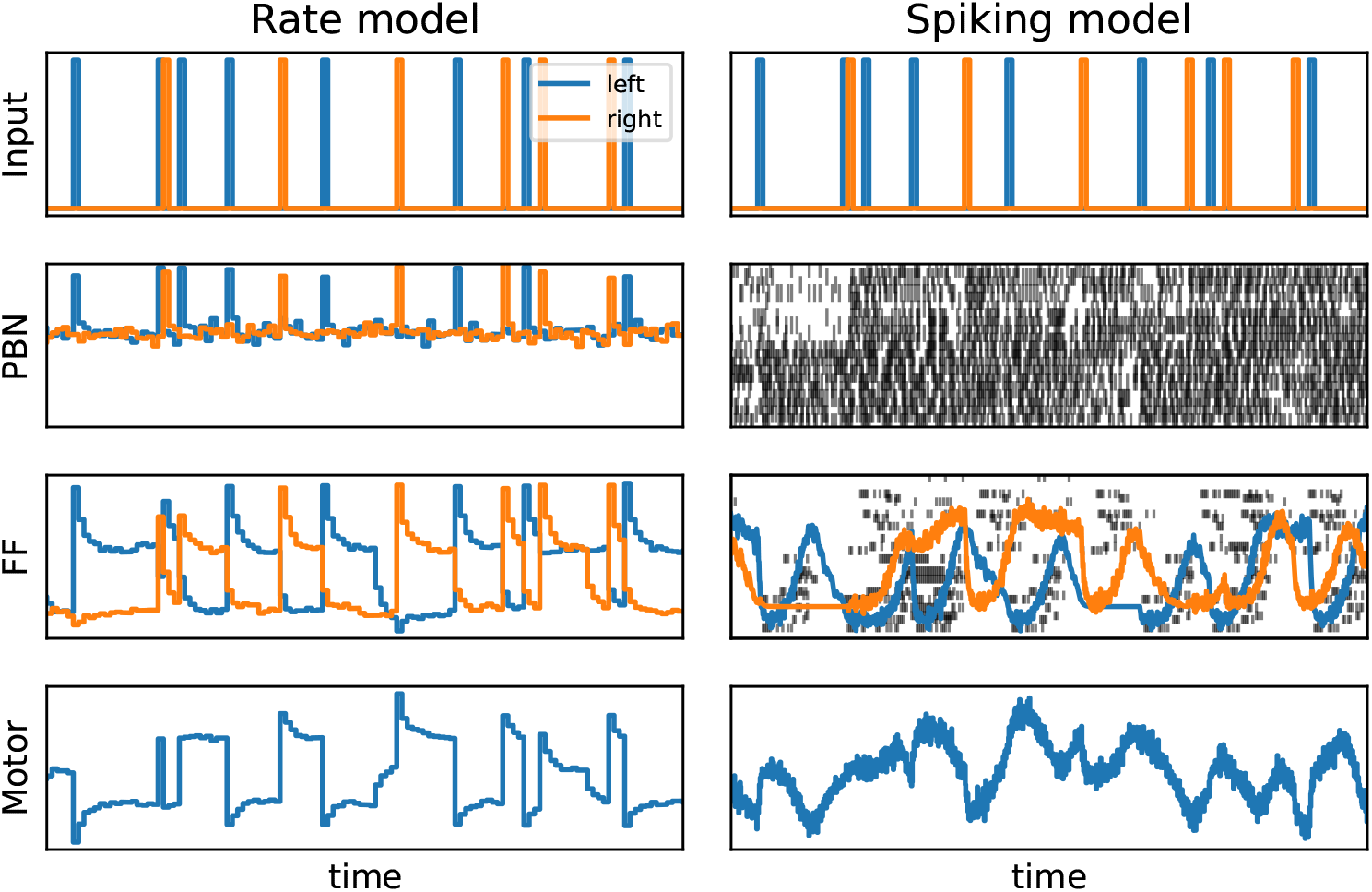
Flip-flopping behaviour of both models.

**S2 Fig.**
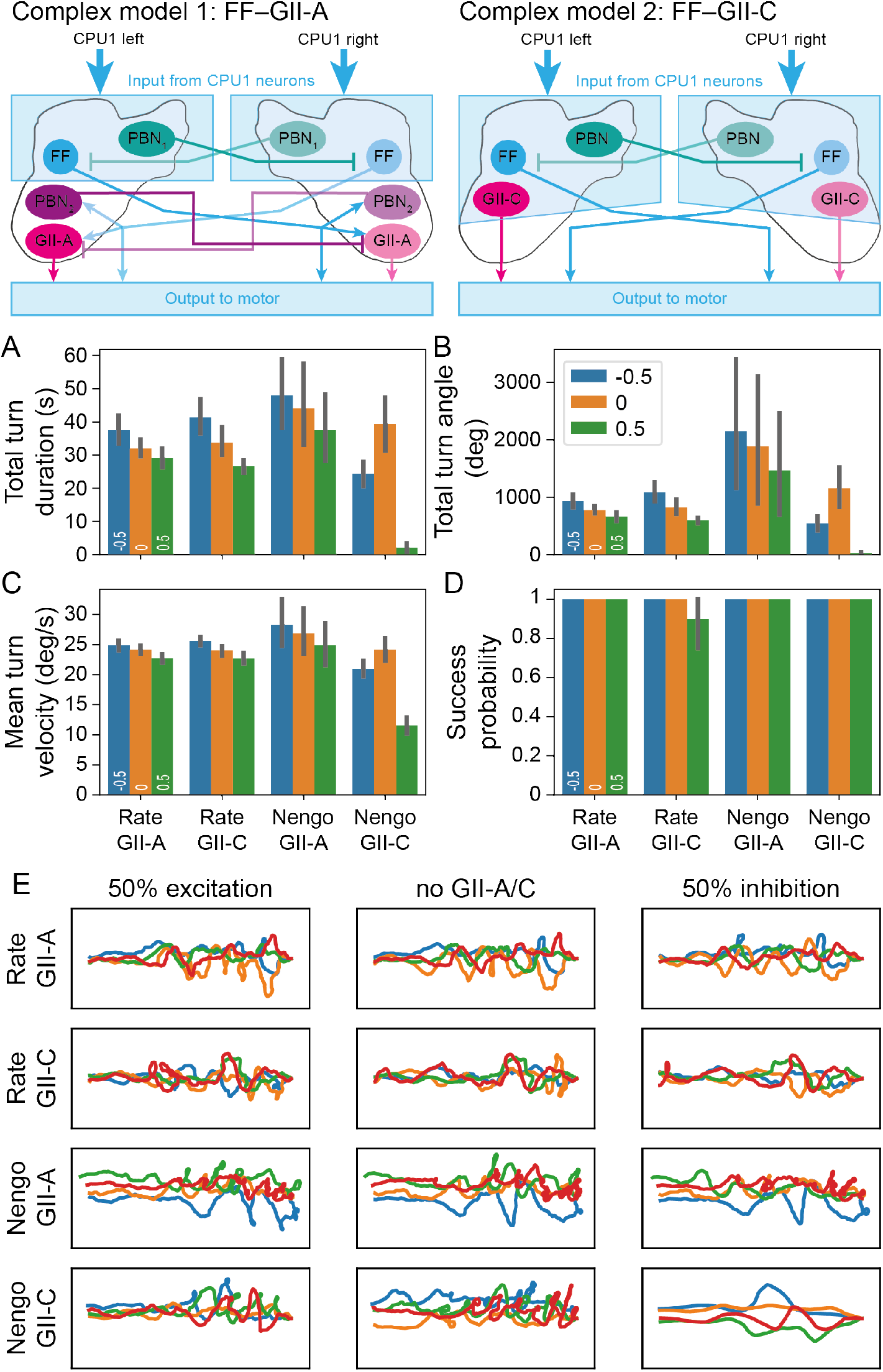
Model results including descending neurons GII-A and GII-C. The GII-A and GII-C cell populations were modelled as inhibitory (−0.5; 50 percent inhibitory input on the motor) or excitatory (0.5; 50 percent excitatory input on the motor).

## Notes

https://github.com/stanleyheinze/insectsteering

### Competing Interest Statement

The authors have declared no competing interest.

